# Testing the reinforcement learning hypothesis of social conformity

**DOI:** 10.1101/2020.03.11.988220

**Authors:** Marie Levorsen, Ayahito Ito, Shinsuke Suzuki, Keise Izuma

## Abstract

Our preferences are influenced by the opinions of others. The past human neuroimaging studies on social conformity have identified a network of brain regions related to social conformity that includes the posterior medial frontal cortex (pMFC), anterior insula, and striatum. It was hypothesized that since these brain regions are also known to play important roles in reinforcement learning (i.e., processing prediction error), social conformity and reinforcement learning have a common neural mechanism. However, these two processes have previously never been directly compared; therefore, the extent to which they shared a common neural mechanism had remained unclear. This study aimed to formally test the hypothesis. The same group of participants (n = 25) performed social conformity and reinforcement learning tasks inside a functional magnetic resonance imaging (fMRI) scanner. Univariate fMRI data analyses revealed activation overlaps in the pMFC and bilateral insula between social conflict and unsigned prediction error and in the striatum between social conflict and signed prediction error. We further conducted multi-voxel pattern analysis (MVPA) for more direct evidence of a shared neural mechanism. MVPA did not reveal any evidence to support the hypothesis in any of these regions but found that activation patterns between social conflict and prediction error in these regions were largely distinct. Taken together, the present study provides no clear evidence of a common neural mechanism between social conformity and reinforcement learning.

## Introduction

Humans are highly sensitive to social influence, and our everyday decisions are often guided by the opinions of others. One of the best known forms of social influence is social conformity, which refers to the act of changing one’s judgments, attitudes, and preferences to align with the expectations of others [1]. The neural mechanism underlying this important social phenomenon has been investigated over the past two decades using functional magnetic resonance imaging (fMRI).

A seminal study by Klucharev et al. [2] found that the posterior medial frontal cortex (pMFC) and ventral striatum play important roles in social conformity. They asked participants to rate the attractiveness of female faces, and after rating each face, the participants were presented with the ratings of the same face by another group of people. They found that the larger the difference between an individual’s rating and the group rating, the higher the activity in the pMFC, while the opposite pattern was found in the ventral striatum (i.e., the closer the individual and group ratings, the higher the activity in the ventral striatum). In a study on social conformity that used transcranial magnetic stimulation, it was shown that the pMFC plays a causal role in preference change [3]. The original fMRI findings have been replicated by several studies (e.g., [4-6]). A recent meta-analysis revealed that the insula and pMFC are consistently positively related to social conflict (i.e., the difference between individual and group opinions) and that the ventral striatum is negatively related to social conflict [7].

As these brain regions (especially the pMFC and striatum) are known to play pivotal roles in reinforcement learning, Klucharev et al. [2] proposed that social conformity and reinforcement learning have a common neural mechanism [8, 9]. It has been reported in several human neuroimaging studies and animal neurophysiology studies that ventral striatum activity tracks reward prediction error (i.e., the difference between expected and actual outcomes) and that the pMFC and insula are involved in processing unsigned prediction error (i.e., the absolute degree of deviation from expectations) (see [10] for a recent meta-analysis of human neuroimaging studies).

In addition to the commonly activated regions reported in human neuroimaging studies, social conformity and reinforcement learning are similar in at least the three ways presented below. First, there is a conceptual similarity between social conformity and reinforcement learning as they are processes that involve the adjustment of the behavior of an individual (e.g., rating or choice) based on received feedback (e.g., group opinion or reward) [9]. Second, the neurotransmitter dopamine is known to play a role in both processes. It is well established that dopamine neurons in the midbrain, which is heavily interconnected to the ventral striatum, signal reward prediction error [11]. Furthermore, a pharmacological study with human participants has shown that social conformity effect is modulated by methylphenidate, which indirectly increases extracellular dopamine levels in the brain [12]. Third, several electroencephalogram (EEG) studies have reported an EEG signal over the pMFC called feedback-related negativity (FRN), which is related to prediction error [13], and several EEG studies on social conformity have reported a similar signal over the pMFC that correlates with the difference between individual and group opinions (e.g., [14-16]).

Although these apparent similarities between social conformity and reinforcement learning have been discussed in literature on social conformity (e.g., [2, 8, 9, 12, 14-16]), to the best of our knowledge, these two processes have never been directly compared to each other. Thus, evidence in support of a common neural mechanism is still insufficient. For example, although meta-analyses on social conformity [7] and reinforcement learning [10] suggest that similar brain regions are involved in these two processes, spatial information is limited in meta-analyses (e.g., see [17]). Therefore, the aim of this study was to rigorously test the reinforcement learning hypothesis of social conformity by asking the same sample of participants to perform social conformity and reinforcement learning tasks inside an fMRI scanner. Furthermore, it is increasingly being recognized that even if there are activation overlaps in the same sample of participants, activation overlaps based on traditional univariate fMRI data analysis cannot be considered strong evidence of a common neural mechanism (e.g., [18]). Therefore, we used multi-voxel pattern analysis (MVPA) to obtain more compelling evidence to support or refute the hypothesis [19].

## Methods

Twenty-nine right-handed female students with no history of psychiatric disorders were recruited from the University of Southampton (mean age = 22.12 years). As in the original study [2], only female participants were recruited for the study as previous research suggests that there are gender differences in neural activity related to attractiveness rating (e.g., [20]). Data from 4 participants were not analyzed for the following reasons: excessive head movement (>3 mm) in 1 participant, strong doubts regarding the social conformity manipulation in 2 participants, and technical problems with the response box in 1 participant (this participant could not complete all the fMRI tasks). The final sample consisted of 25 participants (mean age = 22.1 years). Written consent was obtained from all the participants prior to the experiment, and the study was approved by the University of Southampton ethics committee.

### Stimuli

For the social conformity task, 100 digital color photographs of Caucasian women (aged 18–35) were used as stimuli. The images were taken from the set used in the study by Klucharev et al. [2]. All the women in the photographs were moderately attractive and had a moderate smile.

### Experimental procedure

The experiment consisted of two parts, namely an fMRI session and a behavioral session. Prior to the experiment, participants were given instructions and practice trials on the tasks. During the fMRI session, participants were asked to perform the following two tasks: 1) social conformity task and 2) reinforcement learning task. Both tasks were programmed using Psychtoolbox (http://psychtoolbox.org/) with Matlab software (version 2018b, http://www.mathworks.co.uk). Participants completed two runs of each task (a total of four fMRI runs) and each run consisted of 50 trials. The order of the tasks was counterbalanced across participants.

#### Social Conformity task

For each trial, participants were presented with an image of a female face and a 10-point scale (Figure 1a). The participants were asked to rate the attractiveness of each face on a scale of 1 (least attractive) to 10 (most attractive). Each trial consisted of the following 3 phases: 1) rating phase (no time limit, but participants were encouraged to answer as quickly as they can), 2) highlight of response phase (1–5 seconds, mean = 2 seconds), 3) group rating presentation phase (2 seconds). There was an inter-trial interval (ITI) (1–7 seconds, mean = 2.5 seconds) between trials. The participants were asked to indicate their answers using response button handles (with one handle held in each hand). They used both of their index fingers to move a white cursor through the rating scale (e.g., a right index finger button press moved the cursor 1 point to the right). The white cursor remained invisible until the participant pressed a button, and it appeared below the scale when participants started scrolling. The initial position of the cursor in each trial was randomly determined. The participants were asked to use the cursor location to indicate their chosen rating and to press the right thumb button to select their chosen rating. In the highlight of response phase, the rating chosen by the participant was highlighted with a yellow cursor.

**Figure 1.**
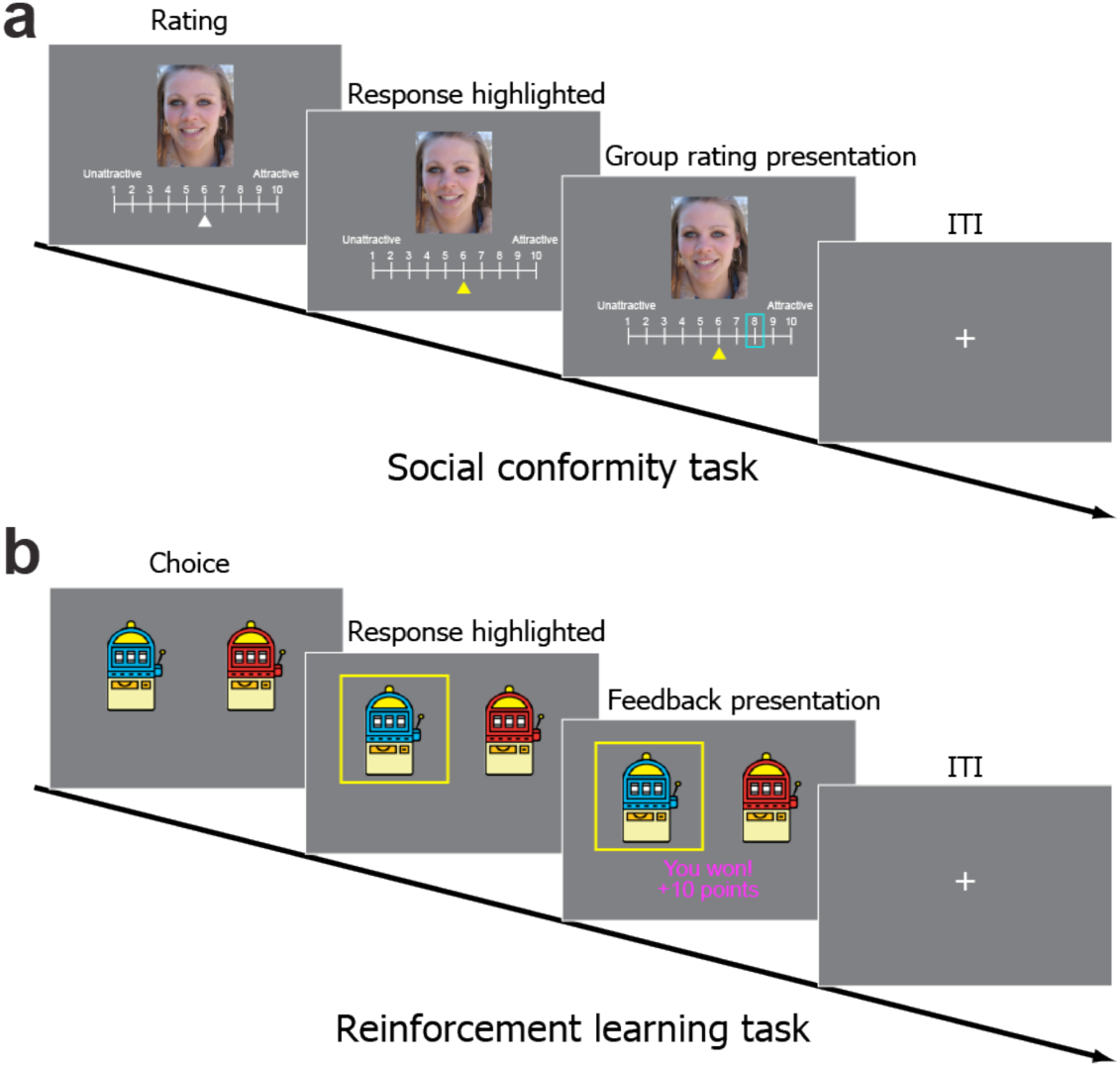
Experimental tasks. **a**) Social conformity task. Participants were shown images of female faces and a 10-point scale and asked to rate the attractiveness of each face. After the participants submitted their ratings, they were shown the ratings of each face by other people (in blue frame) for 2 seconds. **b**) Reinforcement learning task. Participants were presented with two slot machines and asked to pick 1 of them. After the participants made their decisions, they were presented with a win or loss outcome.

In the group rating presentation phase, the participants were presented with a rating in a blue frame that represented “the mean group rating of other students at the University of Southampton” (see Figure 1a). During the instruction, the participants were informed about the meaning of the blue frame and were led to believe that the group rating was an actual group mean rating based on responses from other students at the University of Southampton. In reality, it was systematically manipulated such that the group rating matched the rating of the individual participant in 17.5–25% of the trials and such that the group rating was roughly equally less than or greater than the rating of the participant in 75–82.5% of the trials. The group rating did not deviate from the participant’s rating by more than 3 points. The order of the images was randomized for each participant.

#### Reinforcement Learning task

This was a probabilistic reward learning task adapted from the study by Cooper et al. [21] in which participants were asked to pick one of two slot machines in each trial (Figure 1b). There was a certain probability of winning 10 points (equivalent to £1) on each of the two slot machines. There were independent probabilities of winning on each trial on the two slot machines, and the probabilities changed gradually over the 50 trials in each run to ensure that learning continued throughout the task so that the magnitude of prediction errors varied widely (see [21] for details on how the probabilities were manipulated). There were four different slot machines (red, blue, green, and purple), and two runs of the reinforcement learning task were performed with different combinations of two slot machines. The combinations of the slot machines and the location of the two slot machines were counterbalanced across participants.

Each reinforcement learning trial consisted of the following three phases (Figure 1b): 1) choice phase (participant’s response, <2 seconds), 2) highlight of response phase (1–7 seconds, mean = 2.5 seconds), and 3) outcome phase (2 seconds). Trials were separated by an ITI (1–7 seconds, mean = 2.5 seconds). In the choice phase, the participants were asked to choose the slot machine they thought would result in a win outcome within 2 seconds. They were informed that the probability of winning associated with each slot machine might change gradually throughout the experiment. The participants were also told that two trials would be randomly selected (1 from each of the 2 runs) at the end of the experiment and that they would receive a cash bonus depending on the outcome of the two trials.

The participants were asked to indicate their answers by pressing the response handle buttons using the left or right index finger. If they did not respond within 2 seconds, an error message (“Too slow!!!”) was shown, and the trial was repeated. In the highlight of response phase, the chosen slot machine image was highlighted by a yellow frame. In the outcome phase, 1 of the 2 following possible outcomes was shown: “You won! +10 points” or “You lost. 0 points” (see Figure 1b). The outcome messages were written in magenta or cyan font color, and combinations of font colors (magenta or cyan) and outcomes (win or loss) were counterbalanced across participants.

### Behavioral session

After the scan, to measure how participants’ face ratings were affected by group opinion (i.e., social conformity effect), they were unexpectedly (unannounced during the initial instruction) asked to each face again, this time, without the group rating. They rated the same 100 faces again in a new randomized order. Next, the participants were asked to complete a demographic questionnaire. On the questionnaire, participants were asked to indicate with a “yes” or a “no” if they had any doubts about the group ratings presented during the fMRI face-rating task. If the indication is a “yes,” the participant is asked to explain the doubt in a follow-up interview. As earlier mentioned, two participants were excluded for their strong doubts about the group rating (both of them explicitly said that they did not believe the group ratings presented to them during the fMRI scanning). Lastly, the participants were paid and debriefed.

### fMRI data acquisition

All the images were obtained using a Siemens 3.0 Tesla Skyra scanner. For functional imaging in both task sessions, T2*-weighted gradient-echo echo-planar imaging (EPI) sequences were used with the following parameters: time repetition = 2500 ms, echo time = 25 ms, flip angle = 90°, field of view = 220 mm, and voxel dimension = 3.0 × 3.0 × 3.0 mm. Forty-four contiguous slices with a thickness of 3 mm were acquired in an interleaved order. A high-resolution anatomical T1-weighted image (1 mm isotropic resolution) was also acquired for each participant.

### fMRI data preprocessing

Analysis of the fMRI data was performed using SPM12 (Welcome Department of Imaging Neuroscience) implemented in Matlab (Math Works). To allow for T1 equilibration, the first four volumes were discarded before preprocessing and data analysis. The SPM12 realignment program was used to correct for head motion. Following realignment, the volumes were normalized to MNI space using a transformation matrix obtained from the normalization of the first EPI image of each individual participant to the EPI template using an affine transformation (resliced to a voxel size of 2.0 × 2.0 × 2.0 mm). The normalized data was spatially smoothed with an isotropic Gaussian kernel of 8 mm (full-width at half-maximum). For MVPA, spatial smoothing was not applied so as to preserve fine-grained activation patterns.

### Behavioral analysis

#### Social conformity effect

Multiple regression analysis was performed for each participant to investigate the effect of group rating on individual conformity (rating change). The following two predictor variables were included: 1) gap (group rating - participant’s first rating) and 2) participant’s first rating. The dependent variable was rating change (participant’s second rating - first rating). The first rating was considered as one of the predictor variables to control for the regression-to-the-mean effect [4, 5]. On the rare occasion (0.32% across all participants), seven participants pressed the decide button (right thump button) before the right or left key (i.e., they submitted their rating, most likely accidentally, when the cursor was still invisible). These missed trials were not included in the behavioral data analysis and were modeled as a regressor of no interest in the fMRI data analysis (see below).

#### Prediction error estimation

To estimate prediction error signals in each trial of the reinforcement learning task, we fitted a standard *Q*-learning model [22] to the participants’ choice behaviors.

In the *Q*-learning model, in the choice phase of each trial, an agent chose an option (say *A*) over the other (say *B*) with the probability *q*(A) = 1/[1 + exp(-*β* (*Q*(A) – *Q*(B)))], where *Q* denotes the value of each option and *β* denotes the degree of stochasticity in the choices (called inverse temperature). In the outcome phase, the agent updated the value of the chosen option based on reward experience. Suppose that the option *A* is chosen, then the value is updated by the reward prediction error *δ* = *R* – *Q*(*A*), where *R* denotes the reward outcome (coded 1 for reward and 0 for no reward) as follows: *Q*(A) ← *Q*(A) + *αδ*. Here, the parameter *α* is the learning rate. In the first trial of the task, option values were set at 0.5 (as the agent seemed to have no prior belief in the reward probabilities).

We fitted this model to each participant’s choice data. In the model fitting, to avoid any unreasonable individual fits [23], we employed a maximum a posteriori (MAP) approach in which the learning rate was constrained to a range of 0 to 1 with a Beta (2,2) prior distribution and the inverse temperature was constrained to be positive with a Gamma (2,3) prior distribution.

### fMRI data analysis: univariate analysis

Two general linear models (GLMs) were used to analyze the fMRI data. The first GLM was set up to assess brain activation correlated to the absolute gap (the difference between a participant’s first rating and the group rating) in the social conformity task. The second GLM was set up to assess brain activation correlated to signed and unsigned prediction error values in the reinforcement learning task.

In the first GLM (social conformity task analysis), the absolute gap in the social conformity task was quantified by calculating the absolute difference between the individual and group ratings in each trial. A parametric modulation analysis was performed to assess the correlation between trial-by-trial absolute gap scores and brain activation. The model included the following 3 regressors: 1) trial regressor (onset = trial onset, duration = subject’s response time), 2) feedback regressor (onset = feedback onset, duration = 2 seconds), and 3) feedback regressor modulated by absolute gap between individual and group ratings. As stated above, missed trials were modeled as an additional regressor of no interest for the seven participants.

In the second GLM (reinforcement learning task analysis), signed and unsigned prediction errors were quantified using the computational model described above. A parametric modulation analysis was performed to assess the correlation between trial-by-trial signed/unsigned prediction error and brain activation. The model included the following 4 regressors: 1) trial regressor (onset = trial onset, duration = subject’s response time), 2) feedback regressor (onset = feedback onset, duration = 2 seconds), 3) feedback regressor modulated by signed prediction error values, and 4) feedback regressor modulated by unsigned prediction error values. If there were missed trials, they were separately modeled as a regressor of no interest.

In both GLMs, the regressors were calculated using a box-car function convolved with a hemodynamic-response function. Other regressors of no interest, such as six motion parameters, session effect, and high-pass filtering (128 seconds), were also included.

We aimed to follow up findings based on the univariate analyses (i.e., activation overlaps) with MVPA to further test the hypothesis. To avoid false negative results at the initial univariate analysis stage, we used a statistical threshold of *p* < 0.005 voxelwise (uncorrected for multiple comparisons) with a cluster size of 20 voxels within the three anatomical regions of interest (ROIs, see below) [24]. Outside the ROIs, the statistical threshold was set at *p* < 0.001 voxelwise (uncorrected) and cluster p < 0.05 (FWE corrected for multiple comparisons). All reported *p*-values for both behavioral and fMRI data analyses were based on one-tailed tests.

### fMRI data analysis: MVPA

#### Correlation-based MVPA

When the univariate analyses revealed activation overlaps, we further investigated if activation patterns in each of the overlapped regions were similar between social conflict and signed/unsigned prediction errors (see Figure 2a). In this correlation-based MVPA, the following 4 contrast images were used: 1) those representing positive sensitivity to absolute gaps between individual and group ratings, 2) those representing negative sensitivity to absolute gaps between individual and group ratings, 3) those representing positive sensitivity to unsigned prediction error values, and 4) those representing positive sensitivity to signed prediction error values. We calculated voxel-by-voxel correlations between contrast images 1 and 3 (representing positive social conflict and unsigned prediction error, respectively) for each of the overlapped regions within the pMFC and insula. Similarly, we calculated voxel-by-voxel correlations between contrast images 2 and 4 (representing negative social conflict and signed prediction error, respectively) for each of the overlapped regions within the striatum. These within-subject correlation values were Fisher-z-transformed and submitted to a one-sample t-test to test for significantly positive correlation. A positive correlation indicates that the pattern of voxelwise sensitivity to social conflict is similar to the pattern of voxelwise sensitivity to signed and unsigned prediction errors, thus providing a support for the hypothesis.

**Figure 2.**
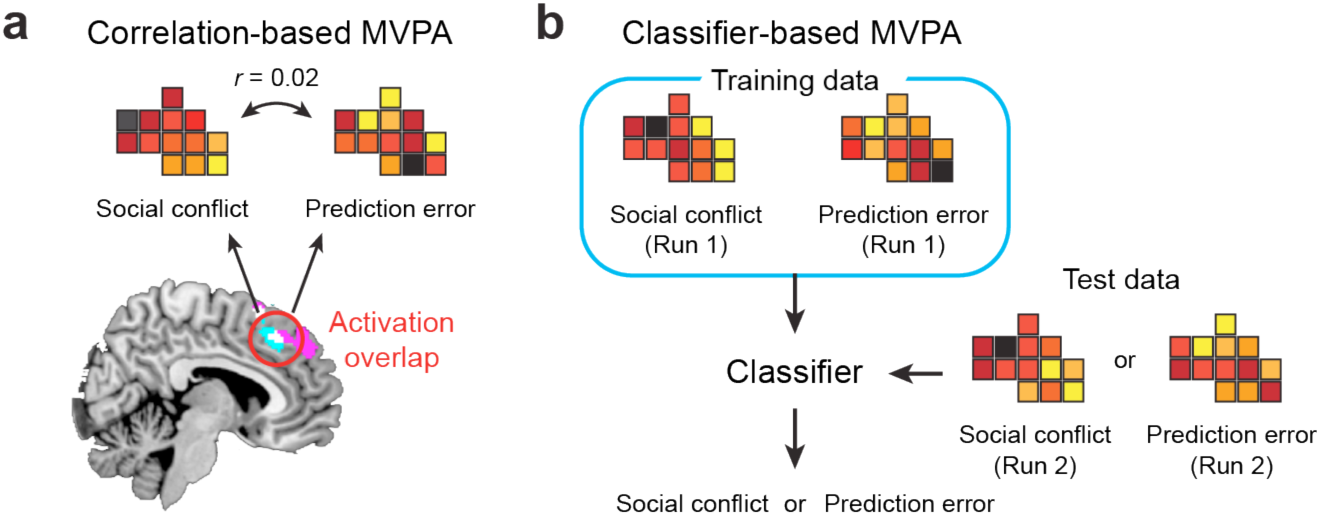
Schematic illustrations of two types of MVPA. **a)** Correlation-based MVPA tests if the two patterns are significantly similar. **b)** Classifier-based MVPA tests if the two patterns are significantly distinct.

#### Classifier-based MVPA

While the correlation-based MVPA described above assessed the similarities in activation (or sensitivity) between social conflict and signed/unsigned prediction errors, the aim of classifier-based MVPA was to determine if the two patterns were significantly distinct. We used a linear support vector machine, which was performed using Matlab in combination with LIBSVM (https://www.csie.ntu.edu.tw/~cjlin/libsvm/) [25], with a cost parameter of c = 1 (default). For this analysis, separate contrast images were created for each of the 2 fMRI runs, and classification performances were evaluated using a leave-one-run-out cross-validation procedure. Thus, using the contrast images from the first run of each task, we trained a classifier that discriminates activation patterns between social conflict and signed/unsigned prediction error. Then, using the contrast images from the second run of each task, we tested if the classifier could discriminate between social conflict and signed/unsigned prediction error (see Figure 2b). The procedure was repeated using data from the second run as training data and data from the first run as test data. Two classification accuracy values were averaged for each participant and the average classification accuracy values were submitted to a Wilcoxon signed-rank test to determine if the classification accuracy was significantly higher than the theoretical chance level (i.e., 50%; note that we also conducted permutation tests [1,000 times] to estimate the empirical chance level in each ROI, but the results were virtually the same). Significantly high classification accuracy indicates that the pattern of voxelwise sensitivity to social conflict is distinct from that to signed/unsigned prediction error.

#### Searchlight analysis

Further, we performed searchlight analysis [26] to more thoroughly depict the activation profiles of each local region within the pMFC, insula, and striatum using the correlation-based and classifier-based MVPA procedures described above. We used a radius of 3 voxels so that each searchlight included a maximum of 123 voxels (and less voxels at the boundaries of each ROI). In each searchlight, a correlation between social conflict and unsigned/signed prediction error was computed for the correlation-based MVPA, and classification accuracy was computed for the classifer-based MVPA. The correlation maps and classification accuracy maps were entered into a second-level permutation-based analysis with 5000 permutations using the Statistical NonParametric Mapping toolbox for SPM [27]. A statistical threshold (i.e., voxel level) was set at *p* < 0.005 and a cluster-level threshold was set at *p* < 0.0125 (FWE corrected; separate second-level analyses were conducted for each of the four ROIs so that the cluster-level threshold was further corrected for four comparisons).

### Regions of Interest (ROIs)

#### Anatomical ROIs

To test the hypothesis, we focused on the following 3 anatomical ROIs: 1) pMFC, 2) insula, and 3) striatum. In a recent meta-analysis, it was reported that these ROIs were consistently positively associated with social conflict (pMFC and insula) and consistently negatively associated with social conflict (striatum) [7]. These ROIs were also consistently associated with signed/unsigned prediction errors [10]. These anatomical ROIs were defined using a WFU pickatlas toolbox for SPM (dilation factor = 2) [28]. The pMFC included the superior frontal gyrus (Frontal_Sup_Medial), anterior cingulate cortex (ACC), middle cingulate cortex, and supplementary motor area (SMA). The striatum ROI included the caudate nucleus, putamen, and globus pallidus. We tested if the same areas within the pMFC and insula ROIs were activated by social conflict (positive correlation) and unsigned prediction error. Similarly, we tested if the same areas within the striatum ROI were activated by social conflict (negative correlation) and signed prediction error.

#### Functional ROIs

For the subsequent MVPAs, we defined functional ROIs as overlapped clusters between social conflict and unsigned prediction error in the pMFC and insula anatomical ROIs and overlapped clusters between social conflict and signed prediction error in the striatum anatomical ROIs with a threshold of p < 0.005 with more than 20 voxels. For the searchlight MVPA, we used the same definition of functional ROIs above but with a more lenient threshold of p < 0.05 (uncorrected).

## Results

### Behavioral results

On average, participants took 5.14 seconds (standard deviation [SD] = 1.65) to rate a face in the social conformity task. During the reinforcement learning task, participants selected a slot machine within 2 seconds for most trials (average number of missed trials = 0.2) and the average reaction time was 0.67 seconds (SD = 0.18).

Consistent with the previous works, we found a significant conformity effect; the second ratings of the participants were significantly influenced by the group rating even after the regression-to-the-mean effect was controlled (*t*(24) = 2.18, *p* = 0.02, *d* = 0.43). We also found highly significant regression-to-the-mean effect (*t*(24) = -15.81, *p* < 0.001, *d* = 3.16), which is consistent with our previous studies [4, 5].

During the reinforcement learning task, participants selected options with a higher reward probability 59.9 % of the trials, which is significantly higher than the chance (50%; *t*(24) = 6.22, *p* < 0.001, *d* = 1.24), indicating that the participants were generally able to accurately keep track of the fluctuating reward probabilities based on reward outcomes.

Further, our data showed that the reinforcement learning model explains participant behavior better than a model that assumes that an individual selects the right option with a fixed probability (*p*). To compare the goodness-of-fit of the models, we computed the Laplace approximated log model evidence [29] of the two models. The values were compared using the Bayesian Model Selection (BMS) method from the study by Stephan et al. [30], which treats model identity as a random effect. Exceedance probabilities from this analysis indicated that the reinforcement learning model has a 100% chance of being the more common of the two models in the population.

Finally, we computed the across-subject correlation between the social conformity effects (beta values from the multiple regression analyses) and learning rate parameters (*α* estimated using the reinforcement learning model), but they did not correlate with each other (*r*_*s*_(23) = -0.09, *p* = 0.66). This indicates that individual differences in the susceptibility to social influence during the social conformity task is unrelated to individual differences in the sensitivity to reward outcome during the reinforcement learning task.

### Univariate Results

We first successfully replicated the findings of previous studies on social conformity. The pMFC (dmPFC [dorsomedial prefrontal cortex] and pre-SMA [pre-supplementary motor area]) and bilateral anterior insula activities were positively correlated with social conflict (i.e., absolute difference between participant and group ratings) (Figure 3a), whereas the striatum activities were negatively correlated with social conflict (Figure 3b and Table 1). All activated areas outside the ROIs are listed in Supplementary Table 1.

**Table 1.** ROI activation during the social conformity task.

| Location | MNI coordinate |  |  | Z | Cluster size |
| --- | --- | --- | --- | --- | --- |
|  | x | y | z |  |  |
| *Areas in the pMFC and insula positively related to social conflict* |  |  |  |  |  |
| dmPFC | 8 | 48 | 38 | 4.26 | 337 |
| Right anterior insula | 38 | 22 | -4 | 4.07 | 388 |
| Left anterior insula | -34 | 20 | -16 | 3.76 | 128 |
| pre-SMA | 12 | 20 | 68 | 3.48 | 191 |
| *Areas in the striatum negatively related to social conflict* |  |  |  |  |  |
| Left striatum (caudate body) | -18 | -8 | 20 | 5.13 | 1,714 |
| Left caudate tail | -14 | -20 | 22 | 4.55 |  |
| Left putamen | -32 | -16 | 6 | 4.22 |  |
| Left caudate head | -18 | 22 | 14 | 4.18 |  |
| Left NAcc | -18 | 10 | -8 | 4.61 |  |
| Right striatum (caudate head) | 14 | 28 | 0 | 4.85 | 1,682 |
| Right caudate body | 20 | 16 | 20 | 4.53 |  |
| Right putamen | 36 | -8 | -4 | 4.3 |  |
| Right caudate tail | 22 | -24 | 22 | 4.27 |  |
| Right NAcc | 16 | 10 | -12 | 3.99 |  |

pMFC, posterior medial frontal cortex; dmPFC, dorsomedial prefrontal cortex; pre-SMA, pre-supplementary motor area; NAcc, nucleus accumbens

**Figure 3.**
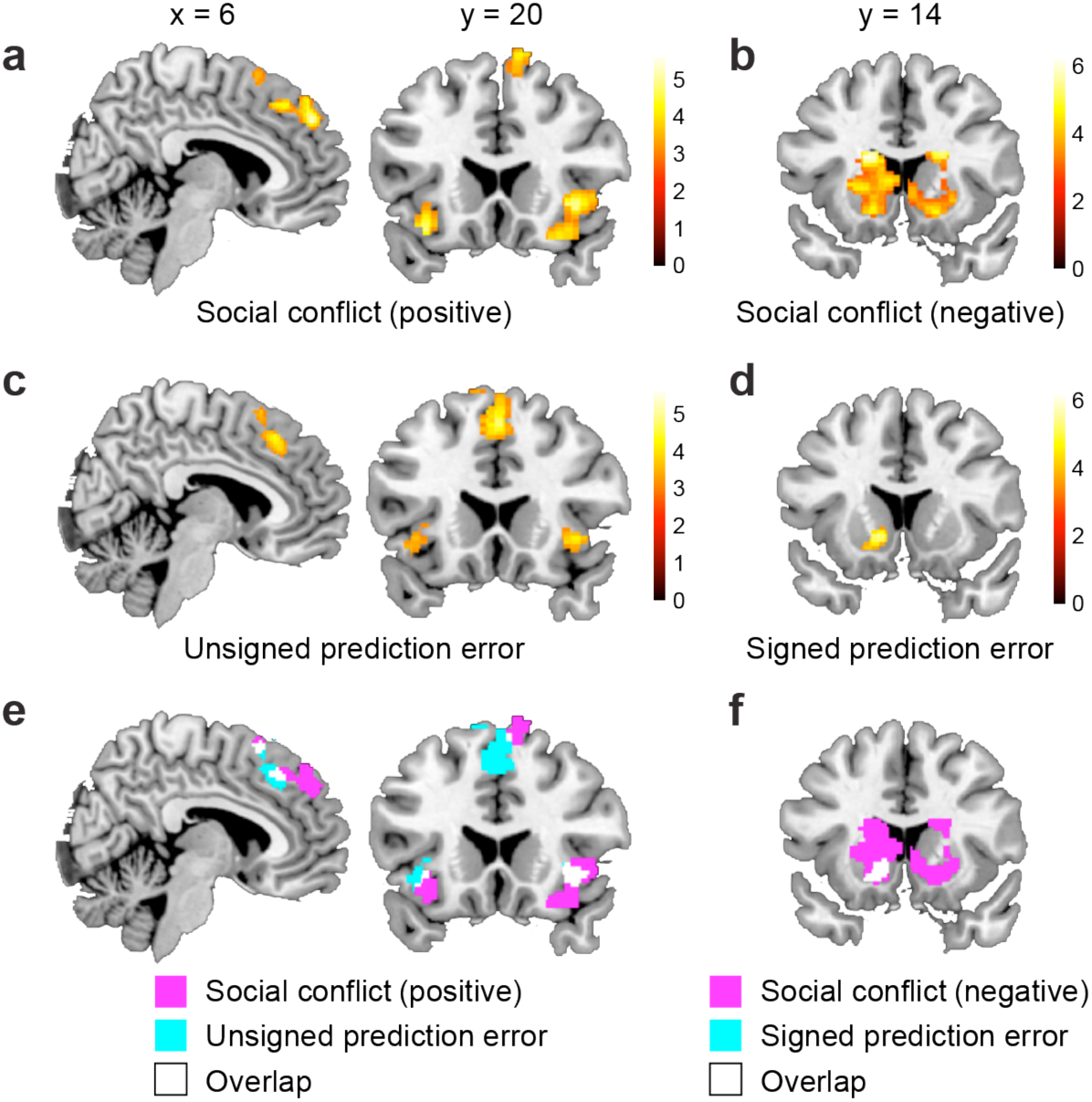
fMRI results from univariate analyses. **a**) pMFC and insula regions positively related to social conflict (i.e., absolute difference between participant and group ratings). **b**) Striatum regions negatively related to social conflict. **c**) pMFC and insula regions positively related to unsigned prediction error. **d**) Striatum regions positively related to signed prediction error. **e**) Activation overlaps between social conflict (panel a) vs. unsigned prediction error (panel c) related regions. **f**) Activation overlaps between social conflict (panel b) vs. unsigned prediction error (panel d) related regions.

We also successfully replicated the findings of previous studies on reinforcement learning. During the reinforcement learning task, the pMFC and bilateral anterior insula activities were positively correlated with unsigned prediction error (Figure 3c), whereas the striatum activities were positively correlated with signed prediction error (Figure 3d and Table 2). All areas outside the ROIs that were significantly positively related to unsigned or signed prediction error are listed in Supplementary Table 2 (note that no area was significantly negatively related to signed or unsigned prediction error).

**Table 2.** ROI activation during the reinforcement learning task.

| Location | MNI coordinate |  |  | Z | Cluster size |
| --- | --- | --- | --- | --- | --- |
|  | x | y | z |  |  |
| *Areas in the pMFC and insula positively related to unsigned prediction error* |  |  |  |  |  |
| mPFC | 16 | 64 | 2 | 4.17 | 55 |
| pre-SMA/dmPFC | -2 | 16 | 54 | 3.99 | 632 |
| Left anterior insula | -44 | 16 | -10 | 3.55 | 89 |
| Right anterior insula | 38 | 18 | -4 | 3.30 | 96 |
| *Areas positively related to signed prediction error in the striatum ROI* |  |  |  |  |  |
| Left putamen | -34 | -4 | 2 | 4.10 | 183 |
| Right putamen | 32 | -6 | 14 | 3.89 | 146 |
| Left NAcc | -12 | 14 | -4 | 3.85 | 224 |

pMFC, posterior medial frontal cortex; mPFC, medial prefrontal cortex; pre-SMA, pre-supplementary motor area; dmPFC, dorsomedial prefrontal cortex; NAcc, nucleus accumbens

Consistent with the hypothesis, in each of the three anatomical ROIs (i.e., the pMFC, insula, and striatum), there were a total of seven activation overlaps (18–158 voxels; see Table 3). We found two overlapped clusters in the pMFC; one in the dmPFC and the other in the pre-SMA (see Figure 3e and Table 3). These overlapped areas in the pMFC and bilateral insula were sensitive to social conflict (positively related) and unsigned prediction error. Similarly, we found three separate overlapped clusters within the striatum (see Figure 3f and Table 3), and these areas were sensitive to social conflict (negatively related) and signed prediction error.

**Table 3.** Overlapped activations between social conflict and signed/unsigned prediction error.

|  | Size of overlap (voxel) | Peak MNI coordinate |  |  |
| --- | --- | --- | --- | --- |
|  |  | x | y | Z |
| *Overlap between areas positively related to social conflict and areas positively related to unsigned prediction error* |  |  |  |  |
| dmPFC | 27 | 8 | 28 | 44 |
| pre-SMA | 26 | 6 | 16 | 62 |
| Right anterior insula | 65 | 38 | 22 | -4 |
| Left anterior insula | 18 | -36 | 20 | -8 |
| *Overlap between areas negatively related to social conflict and areas positively related to signed prediction error* |  |  |  |  |
| Right posterior putamen | 66 | 32 | -12 | 2 |
| Left posterior putamen | 74 | -30 | -12 | 6 |
| Left NAcc | 158 | -18 | 10 | -8 |

dmPFC, dorsomedial prefrontal cortex; pre-SMA, pre-supplementary motor area; NAcc, nucleus accumbens
Note that the peak MNI coordinates reported here are based on the social conflict contrasts. There were two more overlapped clusters (both in the left putamen), but these clusters consist of less than 3 voxels and were not investigated further.

### MVPA Results

#### ROI-based analyses

##### Correlation-based MVPA

To obtain more compelling evidence of a common neural mechanism between social conformity and reinforcement learning, we conducted correlation-based MVPA which investigates whether social conflict and signed/unsigned prediction error evoked similar activation patterns in each of the seven overlapped areas (Table 3). A high correlation indicates that voxels sensitive to social conflict are also sensitive to reward prediction error and therefore supports the hypothesis. However, there were no significant correlations in any of the seven overlapped areas. The average correlations in the four overlapped areas in the pMFC and insula were not significantly positive even at *p* < 0.05 uncorrected level (all *ps* > 0.60; Table 4). Similarly, all three overlapped regions in the striatum showed non-significant correlation (all *ps* > 0.41; see Table 4). These results suggest that, although the same brain regions are involved in social conformity and reinforcement learning, the underlying neural populations may be distinct.

**Table 4.** MVPA results.

| Regions | ROI size (voxel) | Correlation-based MVPA |  | Classifier-based MVPA |  |
| --- | --- | --- | --- | --- | --- |
|  |  | Average correlation | *p*-value (uncorrected) | Average Classification Accuracy (%) | *p*-value (uncorrected) |
| *Overlap between areas positively related to social conflict and areas positively related to unsigned prediction error* |  |  |  |  |  |
| dmPFC | 27 | -0.02 | 0.62 | 59 | 0.004* |
| pre-SMA | 26 | -0.02 | 0.61 | 56 | 0.078 |
| Right insula | 65 | -0.03 | 0.70 | 52 | 0.250 |
| Left insula | 18 | -0.01 | 0.60 | 56 | 0.031 |
| *Overlap between areas negatively related to social conflict and areas positively related to signed prediction error* |  |  |  |  |  |
| Right putamen | 66 | -0.02 | 0.72 | 56 | 0.016* |
| Left putamen | 74 | 0.01 | 0.41 | 51 | 0.500 |
| Left NAcc | 157 | -0.02 | 0.72 | 52 | 0.375 |

\* $p < 0.05$ (Bonferroni correction). Significant results of correlation-based MVPA mean that activation patterns are similar between social conflict and prediction error (i.e., evidence supporting the hypothesis), while those of classifier-based MVPA mean that they are distinct (i.e., evidence against the hypothesis).

##### Classifier-based MVPA

We further attempted to find more direct evidence that refutes the hypothesis and conducted classifer-based MVPA to determine whether activation patterns are distinct between social conflict and signed/unsigned prediction error (i.e., whether a classifer is able to distinguish patterns associated with social conflict and reward prediction error). The dmPFC cluster showed significant classification accuracy, which indicates that the patterns evoked in the dmPFC by social conflict and unsigned prediction error are distinct. Classification accuracies in the right insula, left insula, and pre-SMA were not significant (although classification accuracy in the left insula was significant at *p* < 0.05 uncorrected level; see Table 4). Similarly, the right putamen cluster showed significant classification accuracy, which indicates that the patterns evoked in this area by social conflict and signed prediction error were distinct. Classification accuracies in the left putamen and left nucleus accumbens (NAcc) were not significant (Table 4).

Overall, our ROI based MVPA analysis didn’t find any support for the hypothses. On the contrary, we found evidence refuting the hypothesis, especially in the dmPFC and the right putamen, whereas there was no clear evidence for or against the hypothesis in the pre-SMA, left and right insula, left putamen, and left NAcc.

#### Searchlight analysis

Although we conducted MVPA analyses on each of the seven overlapped clusters (Tables 3 and 4), the size of an overlapped cluster depends on the thresholds and smoothing kernels (e.g., see [31]), and this makes our choice of functional ROIs somewhat arbitrary. Therefore, we conducted searchlight analysis to more thoroughly depict the activation profiles of each local area within the pMFC, insula, and striatum. We defined the functional ROIs (i.e., univariate activation overlaps) more broadly by using a *p* < 0.05 (uncorrected) threshold in each of the three anatomical ROIs. This procedure revealed 917 overlapped voxels in the pMFC, 369 in the right insula, 199 in the left insula, and 3,054 across three separate clusters in the striatum (Figure 4a).

**Figure 4.**
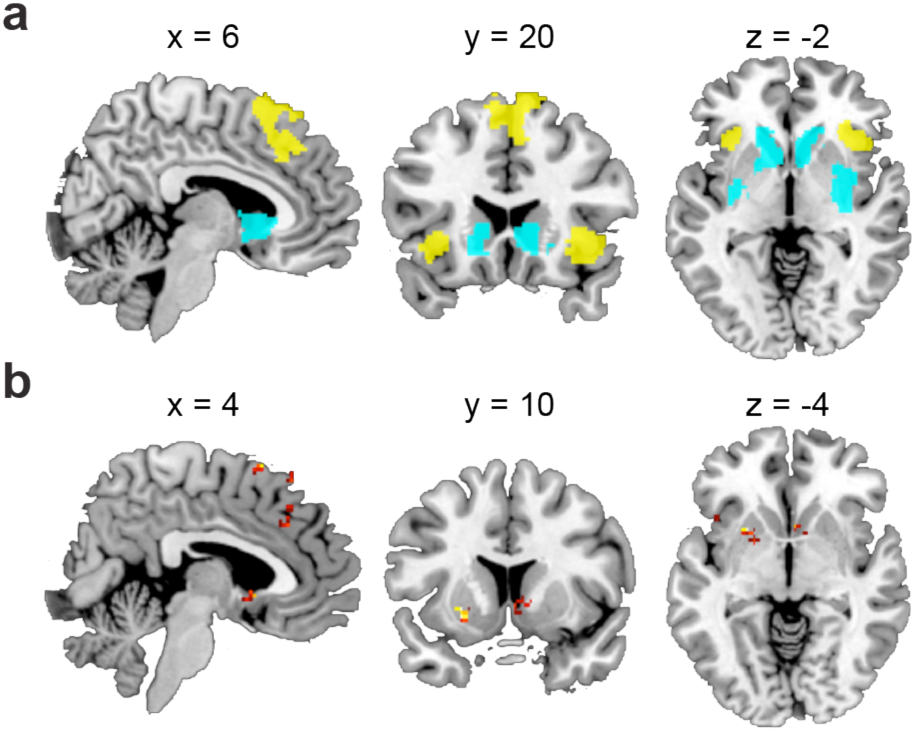
Searchlight MVPA. **a**) Functional ROIs used in the searchlight MVPAs. Yellow color denotes regions sensitive to social conflict (positively related) and unsigned prediction error. Cyan color denotes regions sensitive to social conflict (negatively related) and signed prediction error. **b**) Searchlight MVPA results (classifier-based MVPA). Each panel depicts areas that showed significantly distinct activation patterns between social conflict and signed/unsigned prediction error.

We performed searchlight MVPA of each of the overlapped clusters to determine if any area showed similar activation patterns between social conflict and prediction error (i.e., correlation-based MVPA). However, we found no such areas in any of the ROIs. We also performed classifier-based MVPA using the same searchlight procedure and found a total of nine significant clusters across the four ROIs (Figure 4b and Table 5). Thus, no evidence of a shared neural mechanism between social conformity and reinforcement learning was found in the searchlight analysis. On the contrary, searchlight analysis found that the patterns between social conformity and reinforcement learning in several local areas within each ROI were significantly distinct.

**Table 5.** Searchlight MVPA results.

| Location | MNI coordinate |  |  | Cluster size | Cluster p-value |
| --- | --- | --- | --- | --- | --- |
|  | x | y | z |  |  |
| *Correlation-based MVPA* |  |  |  |  |  |
| No significant region |  |  |  |  |  |
| *Classifier-based MVPA* |  |  |  |  |  |
| pMFC ROI |  |  |  |  |  |
| dmPFC | 16 | 24 | 60 | 54 | < 0.001 |
| dACC | -2 | 30 | 38 | 42 | < 0.001 |
| pre-SMA | 4 | 16 | 66 | 12 | 0.009 |
| Right Insula ROI |  |  |  |  |  |
| Right anterior insula | 44 | 20 | -12 | 5 | 0.001 |
| Left Insula ROI |  |  |  |  |  |
| Left anterior insula 1 | -28 | 24 | 0 | 9 | 0.001 |
| Left anterior insula 2 | -42 | 16 | -4 | 2 | 0.012 |
| Striatum ROI |  |  |  |  |  |
| Left posterior putamen | -24 | 10 | -4 | 29 | < 0.001 |
| Right NAcc | 4 | 12 | -4 | 24 | 0.005 |
| Left anterior putamen | -32 | 2 | 4 | 30 | 0.003 |

## Discussion

The present study investigated whether social conformity and reinforcement learning have a common neural mechanism. The behavioral results showed a robust social conformity effect during the social conformity task and that the reinforcement learning model can adequately explain the behavior of participants during the reinforcement learning task. Furthermore, univariate fMRI analyses successfully replicated the findings of earlier studies that reported the involvement of the pMFC, insula, and striatum in the processing of prediction error signals and social conflict signals during the reinforcement learning task and social conformity task, respectively. Further, we found that, in the pMFC and anterior insula, areas positively related to social conflict (i.e., absolute difference between participant and group ratings) overlapped with areas sensitive to unsigned prediction error. We also found that areas of the striatum negatively related to social conflict overlapped with areas of the striatum sensitive to signed prediction error. This is the first unequivocal evidence that social conflict and signed/unsigned prediction error activate the same areas in the pMFC, insula, and striatum.

However, follow-up MVPA did not provide evidence of a common neural mechanism. It revealed that patterns of sensitivity to social conflict were not similar to patterns of sensitivity to unsigned/signed prediction error in any of the overlapped areas. Although this negative result could be explained by the high noise of the data, classifier-based MVPA could successfully distinguish between social conflict vs. unsigned prediction error related activation patterns in the pMFC, and this is evidence of distinct (non-similar) neural mechanisms in social conformity and reinforcement learning at least within the pMFC. Searchlight analyses further confirmed these results, and there was no evidence of a common neural mechanism in any of the ROIs.

On the contrary, overall pictures of the searchlight results show largely distinct, rather than common, activation patterns in social conflict and signed/unsigned prediction error. Thus, despite our efforts to minimize the possibility of false negatives, we found no evidence to support the hypothesis that social conformity and reinforcement learning have a common neural mechanism. These results suggest that the reinforcement learning hypothesis may be too simplistic to explain the neural mechanism of social conformity.

One possible explanation is that while activities in the pMFC and insula may reflect the degree of surprise (or the absolute deviation from expectations) during the reinforcement learning task, they may express a negative feeling (e.g., unpleasantness or anxiety due to the recognition that one is different from others) during the social conformity task. This in turn motivates the individual to reduce the negative feeling by conforming to group opinion. It is unlikely that individuals think their rating will always be the same as the group rating. In other words, social conflict is not the same as surprise. In fact, using a similar social conformity paradigm, we had previously asked participants to guess the group ratings, but their expectations of group ratings were not related to their own ratings [5]. This indicates that the degree of social conflict is not related to the degree of surprise (deviation from expectations). Furthermore, the notion that the activities in the pMFC and insula during the social conformity task reflect negative emotion is consistent with reports of previous studies stating that these regions also play a role in cognitive dissonance [32, 33] (for reviews, see [8, 34]), which is considered to be a negative feeling caused by inconsistency between attitude and behavior [35]. Nonetheless, negative emotion is typically associated with a more ventral part of the pMFC (i.e., the dACC, rather than the dmPFC or the pre-SMA) [36], although it can be argued that the neural bases of a complex self-conscious emotion like cognitive dissonance may be different from those of more basic emotions such as anger and disgust. This notion should be investigated in future studies.

Similarly, activity in the striatum, which is negatively related to social conflict, may reflect a positive subjective feeling [6, 37], which results from the realization that the group has the same opinion as the participant. Further, activity related to signed prediction error reflects a learning signal but not a positive subjective feeling. It is well known that striatum activity tracks subjective pleasantness induced by various stimuli such as faces and foods (e.g., [33, 38]). However, it should also be noted that the ventromedial prefrontal cortex (vmPFC) is also known to be robustly related to subjective pleasantness (e.g., [38-41]), but we did not find any activation negatively related to social conflict in the vmPFC.

Although the present study used an experimental paradigm similar to that of the social conformity study that originally proposed the reinforcement learning hypothesis of social conformity [2], it should be noted that social conformity is not a unitary phenomenon and that some forms of social conformity may be more similar to reinforcement learning. At least three types of motivation for attitude change based on social influence have been identified in psychological studies [1, 42], and they include the following: 1) motivation to be accurate, 2) motivation to obtain social approval from others, and 3) motivation to maintain a positive self-concept (which includes attitude change following cognitive dissonance). The social conformity effect found in the face-rating paradigm used in this study can be explained by the motivation to obtain social approval and/or the motivation to maintain a positive self-concept, but it cannot be explained by the motivation to be accurate as there is no right or wrong answer in facial attractiveness rating. In a situation where individuals strongly believe that group opinion is more accurate than their own opinion (e.g., group opinion ≈ correct performance feedback), social conflict can serve as a teaching signal just like prediction error in the reinforcement learning task. In fact, the pMFC, insula, and striatum are related to prediction error in a semantic learning paradigm where no reward is involved (i.e., acquiring new semantic knowledge based on performance feedback [correct or incorrect]) [43]. In this study, prediction error in each trial was calculated based on a subjective rating of confidence and on feedback received by participants (e.g., a positive prediction error signal is generated when individuals were not confident about their answer but received a correct feedback). Further, activities in the pMFC and insula were found to be positively related to unsigned prediction error, whereas striatum activities were found to be positively related to signed prediction error [43]. Thus, how these brain regions process social conflict might be more similar to reinforcement learning in a different social conformity paradigm where a right answer can be objectively defined (i.e., where social conflict can serve as a strong teaching signal).

Finally, this study highlights the importance of comparing two tasks (cognitive processes) with the same sample of participants and the utility of the multivariate approach in interpreting univariate activation overlaps [19]. Although previous EEG studies [14-16] report the finding of a signal over the pMFC during the social conformity task that resembles the FRN signal found in the reinforcement learning task, it is important to compare these signals with the same participants to clearly determine if they are similar in terms of spatial location and timing.

In conclusion, this study investigated the reinforcement learning hypothesis of social conformity, which states that social conformity and reinforcement learning have a common neural mechanism. The pMFC, bilateral anterior insula, and striatum were found to be involved in processing reward prediction error and social conflict, and this is consistent with the reports of previous studies. Although searchlight analysis revealed that some areas in the posterior striatum showed similar activation patterns between prediction error and social conflict, MVPA did not find evidence of a shared neural mechanism in the pMFA and insula. Overall, there was no clear evidence to support the reinforcement learning hypothesis of social conformity.

## Acknowledgments

We thank Christopher Watson, Chris Everitt, Angela Darekar, and other members of the MRI unit at Southampton University Hospital for their help with fMRI data collection. We also thank Dr. Vasily Klucharev for providing us with the face stimulus set and Dr. Ryuta Aoki for helpful comments on the manuscript.

## Supplementary Tables

**Supplementary Table 1:** Activations outside the ROIs during the social conformity task.

| Location | MNI coordinate |  |  | Z | Cluster size |
| --- | --- | --- | --- | --- | --- |
|  | x | y | z |  |  |
| *Areas positively related to social conflict* |  |  |  |  |  |
| no significant region |  |  |  |  |  |
| *Areas negatively related to social conflict* |  |  |  |  |  |
| Left IPL | -58 | -30 | 38 | 5.39 | 6,834 |
| Left paracentral lobule | -12 | -20 | 60 | 4.83 |  |
| Posterior cingulate cortex | 0 | -34 | 42 | 4.66 |  |
| Left lateral prefrontal cortex | -42 | 50 | 12 | 4.88 | 499 |
| Left inferior temporal gyrus | -54 | -58 | -8 | 4.5 | 751 |
| Right cerebellum | 46 | -70 | -44 | 4.2 | 240 |
| Right posterior insula | 38 | -8 | -4 | 4.13 | 362 |
| Right STG | 62 | -18 | -2 | 4.13 |  |
| Left DLPFC | -36 | 34 | -32 | 3.92 | 206 |
| Left MFG | -22 | -24 | 52 | 3.86 | 262 |
| Left STG | -62 | -28 | 4 | 3.51 | 204 |

IPL, inferior parietal lobule; STG, superior temporal gyrus; DLPFC, dorsolateral prefrontal cortex; MFG, middle frontal gyrus; STG, superior temporal gyrus

**Supplementary Table 2.** Activations outside the ROIs during the reinforcement learning task.

| Location | MNI coordinate |  |  | Z | Cluster size |
| --- | --- | --- | --- | --- | --- |
|  | x | y | z |  |  |
| *Areas positively related to unsigned prediction error* |  |  |  |  |  |
| Right IPL | 50 | -40 | 48 | 5.28 | 2,706 |
| Right inferior temporal gyrus | 58 | -30 | -22 | 5.03 | 491 |
| Left lateral prefrontal cortex | -28 | 46 | 10 | 4.95 | 175 |
| Left IPL | -38 | -46 | 42 | 4.45 | 653 |
| Right DLPFC | 46 | 32 | 42 | 4.18 | 505 |
| Right VLPFC | 40 | 58 | -10 | 3.74 |  |
| *Areas positively related to signed prediction error* |  |  |  |  |  |
| mPFC | 6 | 62 | 8 | 4.55 | 2,420 |
| dmPFC | -14 | 42 | 54 | 4.39 |  |
| ACC | -2 | 44 | 2 | 4.24 |  |
| vmPFC | -6 | 48 | -14 | 3.41 |  |
| Left lingual gyrus | -20 | -90 | 6 | 4.52 | 424 |
| Right lingual gyrus | 28 | -86 | 10 | 4.39 | 891 |

IPL, inferior parietal lobule; DLPFC, dorsolateral prefrontal cortex; VLPFC, ventrolateral prefrontal cortex; mPFC, medial prefrontal cortex; dmPFC, dorsomedial prefrontal cortex; ACC, anterior cingulate cortex; vmPFC, ventromedial prefrontal cortex
Note that the mPFC cluster positively related to signed prediction error did not overlap with the dmPFC region related to social conflict.

